# Prevalence and diversity of *Helicobacter* species in captive wild carnivores, and their implications for conservation management of endangered species

**DOI:** 10.1101/2024.08.22.609206

**Authors:** Wasimuddin, Gulafsha Khan, Sneha Narayan, Neha Sharma, Basavaraj S. Holeyachi, Abdul Hakeem, Archana Bharadwaj Siva, P. Anuradha Reddy

## Abstract

*Helicobacter* species colonize the gastrointestinal tract of various hosts, and have always been of interest due to their potential zoonotic transmission and impact on health of humans and animals. Comprehensive studies involving wild animals from different locations are lacking, hindering our understanding of their natural host range, prevalence, and genetic diversity. Here we investigated the prevalence and genetic diversity of *Helicobacter* species in thirteen wild carnivore species across twenty-two zoos and rescue centers in India by targeting partial 16S rRNA gene. We used sequences obtained from positive samples for phylogenetic analysis, and further evaluated factors influencing *Helicobacter* prevalence with generalised linear mixed models. We analysed 985 faecal samples of which 286 (29%) tested positive for *Helicobacter*, spanning all the host species included in this study. *Helicobacter* prevalence is strongly related to host species and location, and varied from 7.3% in common leopard to 80% in Indian fox. Phylogenetic analysis identified 59 unique genotypes clustered into 11 groups of three major *Helicobacter* types: enterohepatic, gastric, and unsheathed. Diverse *Helicobacter* species associated with diseases in humans and domestic animals were observed, such as *H. canis*, *H. bilis*, along with several novel strains/ species which require formal classification. Overall, this study highlights the wide occurrence and high genetic diversity of *Helicobacter* species in captive wild carnivores in India. Our findings underscore the need for regular health assessments in captive facilities to monitor *Helicobacter* infections, which could impact the health and management of endangered species. Future research should explore *Helicobacter* presence in biotic and abiotic factors in zoo and free-ranging populations following One Health approaches.

## Introduction

The world, as we know it, is changing and reshaping at an unprecedented scale and speed, bringing the last surviving wildlife populations in close proximity to humans and domestic animals. Following the recent COVID-19 pandemic, there is widespread concern about zoonotic disease risk and transmission of new/ more virulent pathogens from animals - both wild and domestic - to humans. What is even more worrying, and is slowly being accepted as a ticking bomb, is the exposure of endangered wild animals to pathogens from humans and livestock (reverse zoonosis) with serious consequences to their survival as well as public health (Gibb et al., 2020). Deforestation, land use conversion, climate change, illegal wildlife hunting & trade, booming pet markets, international trade & travel routes, are all driving the transmission of pathogens from animals to humans and vice versa.

By definition, zoonotic pathogens can infect multiple hosts, and organisms which can infect several orders of animals, especially wild species, can potentially emerge as new threats (Cleveland et al., 2001). The *Helicobacter* genus is one such group of organisms which can infect a wild range of host species. Over the past three decades, this genus of Gram-negative bacteria has expanded to nearly fifty species, many of which can persistently colonize the gastrointestinal tract of various hosts (Ochoa and Collado, 2021). Waits et al. (2017) found that this genus is highly divergent and may soon be reclassified into several genera. Among its members, the gastric *Helicobacter* species are better studied, primarily due to the important and well-known species *H. pylori*, which infects a large proportion of the human population worldwide. *H. pylori* was identified as a major cause of chronic gastritis and significantly increases the risk of developing peptic ulcers and gastric cancer in humans. There is now growing interest in other gastric and enterohepatic *Helicobacter* species due to their association with several gastrointestinal and systemic diseases, cancers, etc. (reviewed in Ochoa and Collado, 2021). Although many *Helicobacter* species are part of the resident gut microflora in normal or asymptomatic hosts, there is also compelling evidence that their presence is associated with various diseases and neoplasia during times of compromised immunity in both humans and animals (Maurer et al., 2005; Hansen et al., 2011; Castiglioni et al., 2012; Rimbara et al., 2012; Mitchell et al., 2014; Yu et al., 2015; Gill et al., 2016; Herstad et al., 2018; Gilbert et al., 2019; Peng et al., 2021).

*Helicobacter* taxa are recognized as emerging potential pathogens owing to their wide presence and increasing zoonotic potential; however, their natural host range, epidemiological significance, and transmission remain obscure (Goldstein and Solnick, 2003; Mladenova-Hristova et al., 2017). For example, while high prevalence and diversity of *Helicobacter* species in domestic and companion animals have been reported worldwide (Moussa et al., 2021), studies on the prevalence of *Helicobacter* species in wild animals, both free-ranging and captive, are few and sporadic (Eaton et al., 1993; Schröder et al., 1998; Cattoli et al., 2000; Dailidiene et al., 2004; Terio et al., 2005; Eppinger et al., 2006; Tegtmeyer et al., 2013; Terio et al., 2018; Cortez et al., 2021). Furthermore, whereas population-based studies of a single *Helicobacter* species in humans have been used to assess intraspecific genetic variation (Linz et al., 2007); very little is known about the level of genetic variation of *Helicobacter* species from wild animal populations. Hence, screening wild animals for the occurrence and prevalence of different *Helicobacter* species is crucial for gaining insights into their genetic diversity, host range, and eco-epidemiology.

Globally, *ex-situ* management and captive breeding of endangered animals are important components of species conservation, aimed at stabilizing and reestablishing dwindling populations in the wild (Snyder et al., 1996; Ramirez et al., 2006; Ballou et al., 2010). However, drastic environmental changes when animals are brought into captivity can cause radical shifts in their microbiota, creating suitable conditions for the propagation of pathogenic microorganisms (Wasimuddin et al., 2017). This not only affects the survival of captive populations and endangers species management and reintroduction programs but also poses significant health threats, as the zoo environment is highly conducive to the exchange of pathogens. *Helicobacter* species have been associated with diseases in captive populations in a few reports, often with negative health outcomes (Fox et al., 1997). For example, acute gastritis associated with *Helicobacter* species is a major cause of cheetah morbidity and mortality worldwide (Terio et al., 2005). However, comprehensive studies investigating the prevalence and diversity of *Helicobacter* species across a wide range of wild host species in various captive facilities are lacking.

In this study, we examine the prevalence and diversity of *Helicobacter* species in several wild carnivore mammals in zoos and rescue centers across India. India represents an enigma, with a high *Helicobacter* prevalence (80%) in human populations varying according to geographical locations (Misra et al., 2014). Thus, it is essential to assess wild captive populations from different regions of India from a *Helicobacter* perspective. We further discuss the implications of our findings for more efficient management of wild animals in captivity.

## Methodology

### Study area and target species

We collected samples from twenty-two zoos and rescue centers across eight states in India (Fig. 1; Table 1) from thirteen carnivore species - Bengal tiger (*Panthera tigris tigris*), lion (*Panthera leo*), common leopard (*Panthera pardus fusca*), snow leopard (*Panthera uncia*), wild dog/ dhole (*Cuon alpinus*), grey wolf (*Canis lupus*), Himalayan wolf (*Canis himalayansis*), golden jackal (*Canis aureus*), Indian fox (*Vulpes bengalensis*), striped hyena (*Hyaena hyaena*), sloth bear (*Melursus ursinus*), Himalayan black bear (*Ursus thibetanus*), and red panda (*Ailurus fulgens*) between November 2021 to January 2023.

**Fig. 1.**
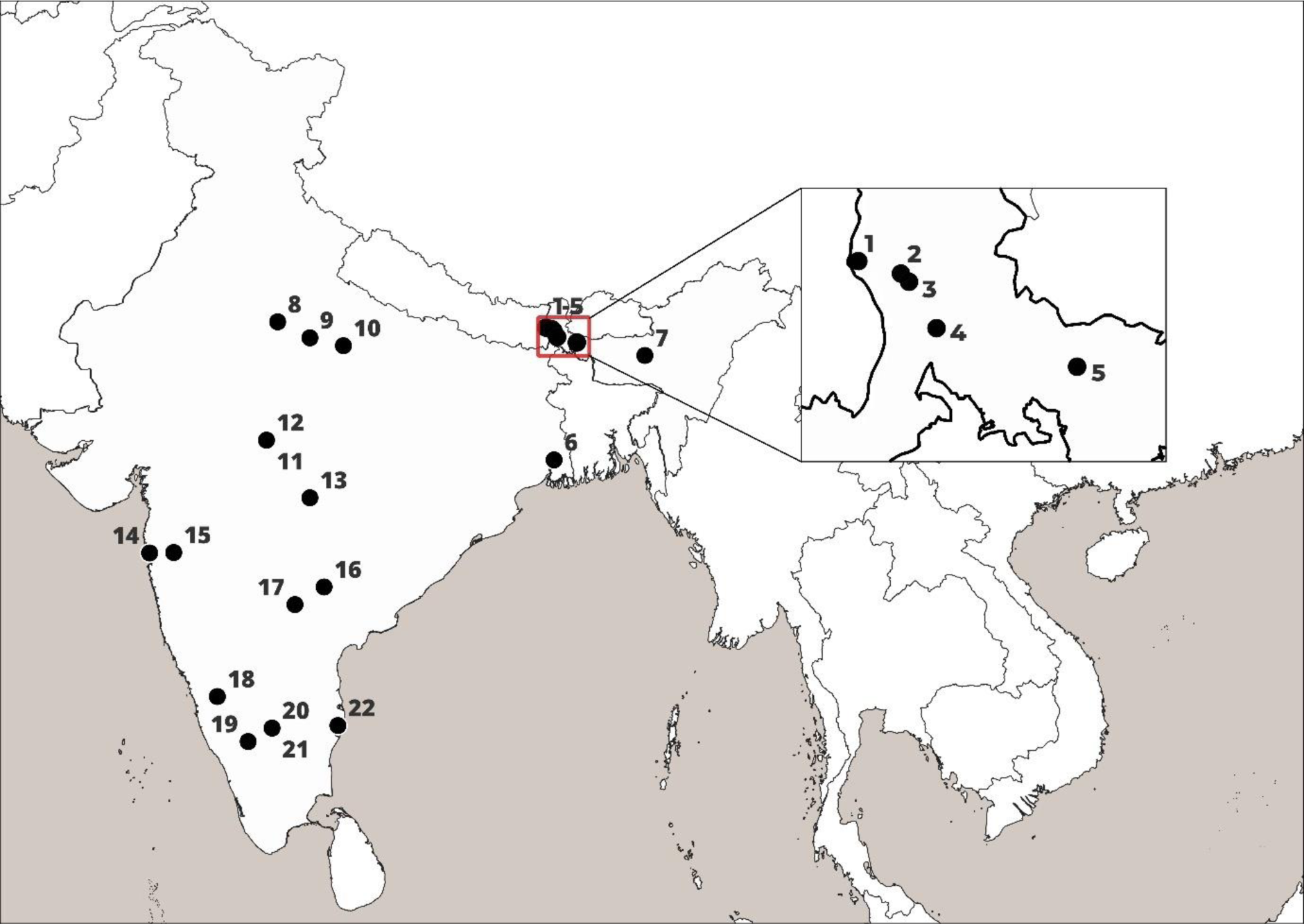
Locations of zoos and rescue centers in India where captive carnivore samples were collected and examined for*Helicobacter* species. 1 – Red panda soft release center, Singhalila National Park (SRC-SNP); 2 – Padmaja Naidu Himalayan Zoological Park, Darjeeling (PNHZP); 3 – Conservation Breeding Center, Tobkedara (NCBC); 4 – North Bengal Wild Animal Park, Siliguri (NBWAP); 5 – Khairbari Tiger Rescue Center, Madarihat (KTRC); 6 – Zoological Garden Alipore, Kolkata (ZGA); 7 – Assam State Zoo, Guwahati (ASZ); 8 – Wildlife SOS, Agra (WSOS-A); 9 – Etawah Lion Safari Park (ELSP); 10 – Kanpur Zoological Park (KNZP); 11 – Van Vihar National Park, Bhopal (VVNP); 12 – Wildlife SOS, Bhopal (WSOS-BH); 13 – Gorewada Rescue Center, Nagpur (GRC); 14 – Sanjay Gandhi National Park, Mumbai (SGNP); 15 – Wildlife SOS, Junnar (WSOS-J); 16 – Kakatiya Zoological Park, Warangal (KAZP); 17 – Nehru Zoological Park, Hyderabad (NZP); 18 – Tiger and Lion Safari, Thevarakoppa (TLST); 19 – Sri Chamarajendra Zoological Garden, Mysuru (SCZG); 20 – Bannerghatta National Park (BNP); 21 – Wildlife SOS, Bannerghatta National Park (WSOS-BNP); 22 – Arignar Anna Zoological Park, Chennai (AAZP).

**Table 1.**
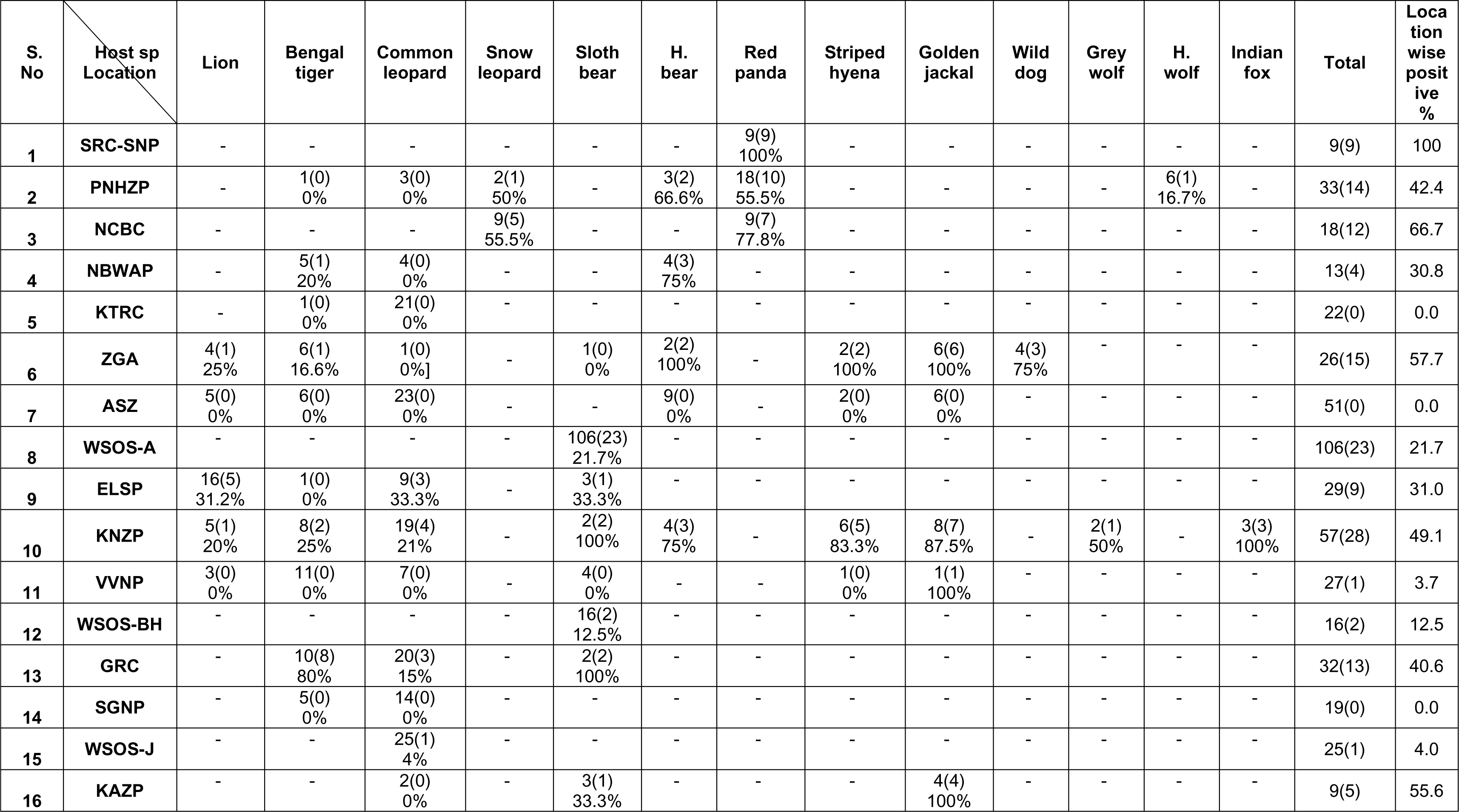
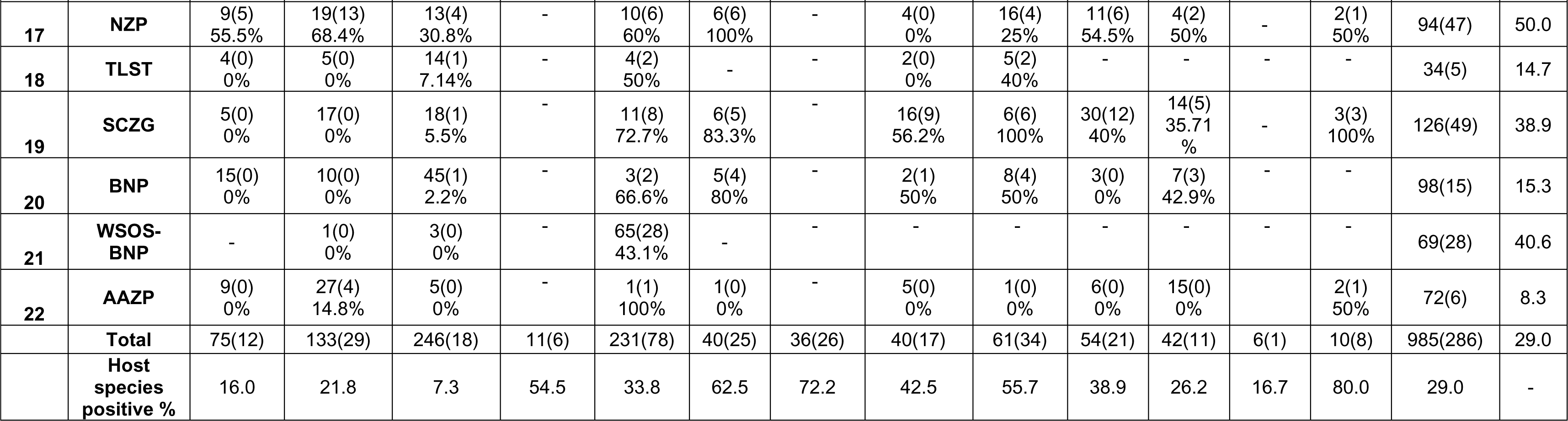
*Helicobacter* prevalence as seen in different host species across various zoos and rescue centers in India. 1 – Red panda soft release center, Singhalila National Park (SRC-SNP); 2 – Padmaja Naidu Himalayan Zoological Park, Darjeeling (PNHZP); 3 – Conservation Breeding Center, Tobkedara (NCBC); 4 – North Bengal Wild Animal Park, Siliguri (NBWAP); 5 – Khairbari Tiger Rescue Center, Madarihat (KTRC); 6 – Zoological Garden Alipore, Kolkata (ZGA); 7 – Assam State Zoo, Guwahati (ASZ); 8 – Wildlife SOS, Agra (WSOS-A); 9 – Etawah Lion Safari Park (ELSP); 10 – Kanpur Zoological Park (KNZP); 11 – Van Vihar National Park, Bhopal (VVNP); 12 – Wildlife SOS, Bhopal (WSOS-BH); 13 – Gorewada Rescue Center, Nagpur (GRC); 14 – Sanjay Gandhi National Park, Mumbai (SGNP); 15 – Wildlife SOS, Junnar (WSOS-J); 16 – Kakatiya Zoological Park, Warangal (KAZP); 17 – Nehru Zoological Park, Hyderabad (NZP); 18 – Tiger and Lion Safari, Thevarakoppa (TLST); 19 – Sri Chamarajendra Zoological Garden, Mysuru (SCZG); 20 – Bannerghatta National Park (BNP); 21 – Wildlife SOS, Bannerghatta National Park (WSOS-BNP); 22 – Arignar Anna Zoological Park, Chennai (AAZP).

### Sample collection and DNA analysis

Fresh faecal samples were collected with a clean spatula, sterilized with bleach and alcohol, within minutes of defecation, after moving the concerned animal to another enclosure with the help of animal keepers. We took several precautions while collecting samples like wearing mask and gloves, avoiding samples’ contact with soil, transferring samples immediately into the sterile tube, which was then sealed properly. Each sample collection tube was labelled with details such as Name/ ID of the animal, gender, location and date of sample collection. Faecal sample tubes were immediately transferred to a -10°C Zedblox Actipod^TM^ to arrest microbial growth and to minimise contamination. DNA was isolated from the faecal samples with Nucleospin DNA stool Kit (Macherey-Nagel, Germany) as per manufacturer’s instructions. We avoided possible contamination by aliquoting samples and isolating DNA in separate biosafety cabinets. DNA quality and quantity was measured using a UV-Nano-drop spectrophotometer. Samples with 20ng/µl of DNA were used directly while those with higher DNA concentrations were diluted to 20ng/µl prior to Polymerase Chain Reaction (PCR). *Helicobacter* genus-specific primers C97F (5’-GCTATGACGGGTATCC-3’) and C98R (5’-GATTTTACCCCTACACCA-3’) targeting ∼398bp segment of 16S rRNA gene (Shen et al., 1997) were used to detect the presence of *Helicobacter* taxa in the faecal samples. A 20μl PCR reaction mixture consisted of 20ng/ul template DNA, 250μM dNTP, 1xTaq buffer (TaKaRa, Japan), 1x BSA, 1U Taq polymerase (TaKaRa Ex Taq Hot Start Version, TaKaRa, Japan), and 5μM of each primer. PCR was set in Veriti^TM^ 96-Well Fast Thermal Cycler (Applied Biosystems, USA) with an initial denaturation at 95°C for 5 minutes followed by 40 cycles of denaturation at 95°C for 30 seconds, annealing at 51°C for 30 seconds, extension at 72°C for 30 seconds and a final extension at 72°C for 5 minutes. Following PCR, the PCR products were visualized in a 2% agarose gel, and purified for Sanger sequencing.

### Sequencing and phylogenetic analysis

PCR amplicons were sequenced in ABI 3730 DNA Analyser (Applied Biosystems, USA). Sequences were trimmed and assembled using CodonCode Aligner (CodonCode Corporation, USA) to generate consensus sequences. We searched for sequence similarities using NCBI-BLASTN (Altschul et al., 1990), and the cut-off for both percentage identity and query coverage was ≥98%. ClustalW multiple sequence alignment was performed in BioEdit (Hall, 1999). A neighbour-joining (NJ) tree (Saitou and Nei, 1987) was constructed in MEGA7 software (Kumar et al., 2016) using K2-P distances among sequences (all trimmed to a length of 323bp) with 1000 bootstraps (Felsenstein, 1985) combining (i) 59 unambiguous sequences from samples in this study (i.e. excluding heterozygous sequences indicating multiple infections with more than one strain or species), (ii) 39 sequences of *Helicobacter* species known to occur in diverse mammalian species (Dewhirst et al., 2005) or (iii) sequences showing high similarity (≥98% identity in BLAST searches) with the sequences obtained within this study. *Wolinella succinogenes* (M88159) and *Campylobacter jejuni* (L04315) were used as outgroups as done previously (Wasimuddin et al., 2012; Ashaolu et al., 2022).

### Statistical analysis

In order to determine if host species and sampling locations can explain *Helicobacter* infection among individuals, we performed generalized linear mixed models (GLMMs) in R environment (R version-4.3.1, R Core Team). We modelled *Helicobacter* infection according to host species (n=13), sampling location (n=22), and sample ID to control the random effect (lme4; Bates et al., 2015). Model selection was based on the information-theoretic (IT) approach using a second-order Akaike’s information criterion corrected for small sample sizes (AICC) and Akaike weights (ω) to determine model support.

## Results

We collected 985 faecal samples from thirteen endangered carnivore species in twenty-two zoos and rescue centers across India. Following PCR and sequence analysis, 286 samples were found to be positive for *Helicobacter*, giving an overall prevalence of 29%. All the studied host species found to be *Helicobacter* positive; however, *Helicobacter* prevalence varied in different species, the lowest being in common leopard (7.3%) and the highest in Indian fox (80%) as shown in Table 1. Location-wise prevalence ranged from 0% in Khairbari Tiger Rescue Center, Assam State Zoo and Sanjay Gandhi National Park; to 100% in Red Panda Soft Release Center, Singhalila National Park (Table 1). Sequence analysis revealed three different major types of *Helicobacter* based on their preferred place of colonization or *Helicobacter* without sheathed flagella; enterohepatic, gastric and unsheathed (Fig. 2). A large proportion (nearly 50%) of the positive samples matched with unclassified *Helicobacter* spp., especially in ursids (sloth bear and Himalayan black bear) and red panda (Table 2). In case of the canids (golden jackal, dhole, Himalayan wolf and Indian fox) and striped hyena, the most frequently matched species were *H. canis*, *H. felis*, *H. cinaedii* and *H. winghamensis* (Table 2).

**Fig. 2.**
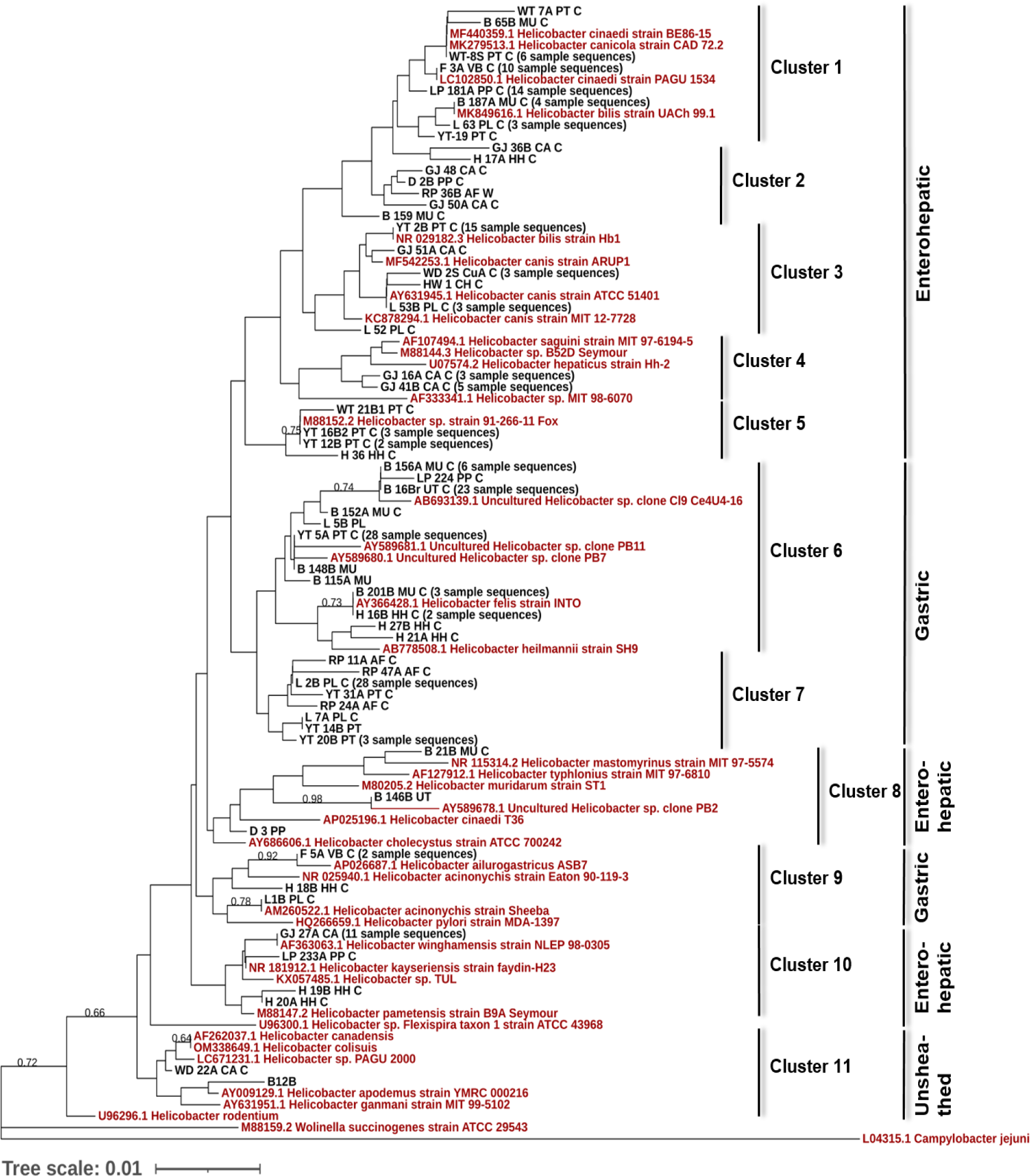
Distribution and evolutionary relationships of *Helicobacter* strains/species in captive carnivore species: Neighbour-joining tree based on comparison of 16S rRNA gene sequences, obtained from this study (59 unique genotypes-in black), sequences of *Helicobacter* strain/species known to occur in diverse mammalian species (in red), and sequences showing high similarity (≥98%) in BLAST searches with the sequences obtained within this study (in red). GenBank accession numbers are shown along with strain/species name. Numbers above branches are bootstrap values (only those above 60 are shown). The sequences are clustered and grouped further based on their preferred place of colonization or if they are unsheathed.

**Table 2.**
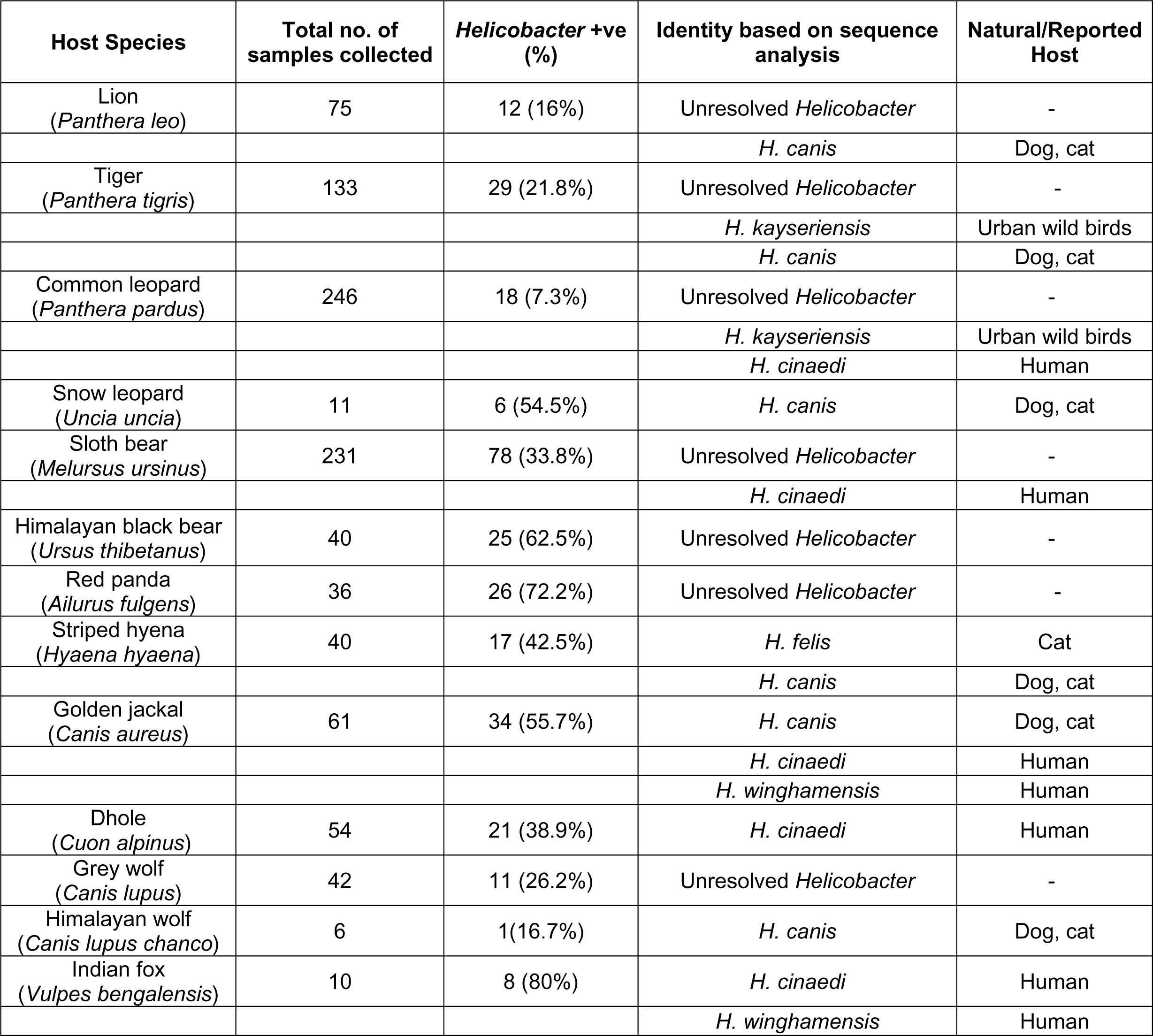
Frequently identified *Helicobacter* species in different studied host species.

We obtained 215 clear sequences from 286 *Helicobacter* positive samples, whereas clean chromatograms could not be obtained from 71 samples due to mixed *Helicobacter* signatures. We found 59 unique genotypes in which 7 major genotypes were noted to represent 129 of 215 sequences (Genbank accession numbers PP961961 – PP962171). The neighbour-joining tree, with 59 genotypes along with 39 reference *Helicobacter* sequences or *Helicobacter* sequences showing close match with sample sequences, showed 11 distinct sequence clusters (Fig. 2) falling into 3 major *Helicobacter* types (i.e. Enterohepatic, Gastric, Unsheathed). Clusters 1-5, 8 and 10 were enterohepatic, whereas clusters 6-7 and 9 were gastric, and cluster 11 was unsheathed *Helicobacter* species. Cluster 6 was the largest cluster representing 69 samples followed by cluster 1 (40 samples), cluster 7 (37 samples), cluster 3 (24 samples), etc. On closer look, cluster 6 was divided into two sub-clusters, one represented by *H. felis* and *H. heilmannii*, and the second largely represented by uncultured *Helicobacter*. Further, cluster 6 mainly consisted of samples from sloth bear (65.22%) and Himalayan black bear (15.94%). Reference *Helicobacter* sequences or sequences with high similarities with our study samples were distributed in all the clusters except cluster 2 and cluster 7, which were found to be unique with no close representatives in the phylogenetic tree (Fig. 2). Cluster 2 consisted of seven unique sequences, three of them originating from golden jackal, and one each from common leopard, sloth bear, hyena and red panda. Cluster 7 consisted of eight unique sequences - three of them originating from red panda, another sequence present in 28 individuals mostly red panda (57%), three sequences from tiger, and one from Asiatic lion. *H. rodentium* (U96296) and *Helicobacter sp*. Flexispira taxon 1 (U96300.1) sequences taken from NCBI as reference sequences, did not fall in any of the clusters.

Generalized linear mixed model showed a strong support for effects of host species and locations in explaining *Helicobacter* prevalence (ΔAIC_C_ = 159.30), by controlling the random effect of sample identity. The conditional coefficient (R^2^GLMM(c)) of determination was 0·601, which explains the variance of both the fixed and random factors. Marginal coefficient (R^2^GLMM(m)) of determination was 0·484, which explains the variance of the fixed factors only, which we calculated as the variance explained by the best model and ΔAIC_C_. Highest *Helicobacter* prevalence was observed in Indian fox (80%) followed by red panda (72.2%), Himalayan black bear (62.5%), golden jackal (55.7%) and snow leopard (54.5%), while the lowest prevalence was noted in common leopard (7.3%) (Table1). In terms of location, the maximum number of *Helicobacter* positive samples were from Red Panda Soft Release Centre, Singhalila National Park (100%) followed by Conservation Breeding Center, Tobkedara (66.7%), Kakatiya Zoological Park, Warangal (55.6%) and Nehru Zoological Park, Hyderabad (50.0%), whereas samples from host species from several locations were found to be *Helicobacter* negative (Table1).

## Discussion

To the best of our knowledge, this study is the most comprehensive screening of diverse wild carnivores housed in various captive facilities across a large and biodiverse region for the presence of *Helicobacter* species. Results of this study indicate that *Helicobacter* is present in all the thirteen wild animal species screened, with an overall prevalence of 29%. However, the prevalence varies among different host species and across various locations. We detected *Helicobacter* presence for first time in some of the host species such as snow leopard, sloth bear, Himalayan black bear, striped hyena, dhole or Indian wild dog, Himalayan wolf and Indian fox. Among the canids, Indian fox and golden jackal show a high prevalence of *Helicobacter* species (Table 1) earlier detected and reported in humans, and domestic dogs and cats (Table 2; Melito et al., 2001; Kaakoush et al., 2010; Kakuta et al., 2014), and both of these host species are commonly present around human settlements. However, mere presence of *Helicobacter* species may not indicate transmissions, especially when these organisms are known to be part of the normal gut flora. Nevertheless, the high prevalence reported here definitely warrants further investigations. Comparatively lower prevalence of *Helicobacter* was observed in lion, tiger and leopard in comparison to other host species, and with reference to an earlier study (Jakob et al., 1997). However, the diversity of species/ strains was high, for example 14 different genotypes were observed in tiger samples, which could be due to high number of samples tested from various locations, compared to other host species.

We observed 59 *Helicobacter* strains/ species from 13 studied captive carnivores, which suggests wide presence and unexplored genetic diversity of *Helicobacter* in wild animals in India. Few *Helicobacter* strains/ species (i.e. 7 unique genotypes) dominate and are present in more than half of the samples sequenced. The detected genotypes showed close sequence similarity with various reference *Helicobacter* species included in the phylogenetic tree. Overall, noted clusters could be broadly divided into three *Helicobacter* types based on either their preferred place of colonization (i.e. enterohepatic, gastric) in the gastrointestinal tract, or *Helicobacter* without sheathed flagella (i.e. unsheathed) as observed in earlier studies (Dewhirst et al., 2005; Ashaolu et al., 2022). In India high occurrence and genetic diversity of *H. pylori* were observed in human populations (Ghosh et al., 2016). Interestingly, only 15-20% of *H. pylori* infected populations in India develop gastric or duodenal ulcers. Although, we did not detect *H. pylori* in our studied samples, high diversity in strains or species could be a cause for concern of the health of captive populations, especially in the case of *Helicobacter* species which are associated with diseases in animals (Eaton et al., 1993; Schröder et al., 1998; Cattoli et al., 2000; Dailidiene et al., 2004; Terio et al., 2005; Eppinger et al., 2006; Tegtmeyer et al., 2013; Terio et al., 2018; Cortez et al., 2021). For example, we observed *H. acinonychis*, associated with cheetah deaths in captivity and also with diseases in other carnivores, in one lion sample (Fig. 2). *Helicobacter acinonychis* has earlier been isolated and characterised from various big cats in different parts of the world (Schröder et al., 1998; Dailidiene et al., 2004; Eppinger et al., 2006; Tegtmeyer et al., 2013), and is found to be closely associated with *H. pylori* (Tegtmeyer et al., 2013). Eppinger et al. (2006) presented compelling genomic evidence of reverse zoonosis and suggested that *H. pylori* jumped from humans to a wild feline host nearly 200 kya (range 100-400 kya), adapted to its new host, speciated as *H. acinonychis* and then spread globally in several feline species like cheetah, tiger and lion. Like *H. pylori* in humans, *H. acinonychis* is associated with chronic gastric pathology in wild felids (Schröder et al., 1998; Cattoli et al., 2000; Tegtmeyer et al., 2013; Mangiaterra et al., 2022).

It should be noted that around 25% of *Helicobacter* positive samples could not be included in our phylogenetic tree due to mixed *Helicobacter* signatures. Therefore, we cannot not rule out the possibility of presence of other strains/ species of *Helicobacter* undermining the overall diversity. The significant effect of sample location on *Helicobacter* prevalence could be largely because samples from different species were collected from different locations. Therefore, in order to tease apart the role of location in shaping *Helicobacter* prevalence, samples from one host species should be collected from different locations. Future research should also account for the effect of other host variables like age, health status, etc. This is usually difficult to achieve as different facilities house various animal species. Nevertheless, we accounted for all available mega- and meso-carnivores from each facility to assess the full breadth of *Helicobacter* presence and its genetic diversity in captive carnivore population in India.

A large proportion of red panda, sloth bear and Himalayan black bear samples showed the presence of *Helicobacter* species which did not match with any of the known or classified *Helicobacter* species or strains (i.e. cluster 6, 7, Fig. 2; Table 2). We obtained similar results in free-ranging red panda samples collected in Neora Valley National Park, West Bengal. This suggests possible presence of novel *Helicobacter* strains or species in these host species, and requires further research and formal classification. An earlier study by Sommer et al. (2016) showed that *Helicobacter* abundance increases in brown bears in summer months in response to diet change after the hibernation period. Interestingly, it has also been observed that cases of *H. pylori-*associated gastritis increase with altitude in human subjects (Sharma et al., 2014; Quiñones-Laveriano et al., 2020), Thus, whether the observed *Helicobacter* species or strains in red panda and Himalayan black bear are opportunistic pathogens or important gut flora associated with cold adaptation or any other vital function in these high-altitude animals remains to be tested. These results also explain why two locations, Red Panda Soft Release Center, Singhalila National Park and Conservation Breeding Center, Tobkedara have the maximum numbers of *Helicobacter* positive samples as these largely or exclusively house red panda. Interestingly, both these locations are fairly isolated, have minimal human footfall, and are dedicated for the conservation of endangered Himalayan species like red panda and snow leopard. Similarly, cluster 2 in our phylogenetic tree has seven genotypes without any close reference *Helicobacter* species, suggesting novel genotypes not recorded earlier. Three of these genotypes are exclusively from golden jackal. All our findings warrant further research in this direction in order to formally classify these genotypes and to ascertain their association with animal health.

## Conclusions

We documented the broad occurrence and high genetic diversity of *Helicobacter* species in wild captive carnivores in India with prevalence varying according to host species. Diverse *Helicobacter* species known to be associated with diseases in humans and domestic animals were observed, but also several novel strains/ species were detected which require further research and formal classification. Our results indicate that *Helicobacter* species could be common residential bacteria in some of the host species, but given their ability to cause severe debilitating disease in multiple hosts, these species should be studied further for their impact on health of wild animals. The zoonotic potential and mode of transmission of these *Helicobacter* species through contaminated food, water or vectors like rats, need to be examined in detail. Future research should examine zoo personnel like animal keepers in order to explore the zoonotic potential of *Helicobacter* species, as well as free ranging wild populations in order to disentangle the role of captive facilities in *Helicobacter* infection following the One-Health framework. We recommend inclusion of potential pathogens such as *Helicobacter* species in regular health assessment in zoos and rescue centers for better health management of animals and their keepers and to record pathogen transmission events, if any.

## Ethics approval

Permissions to collect samples for this study were granted by Chief Wildlife Wardens of West Bengal (No.1703/WL/4R-31/2021, dated 01/09/2021), Maharashtra (No: Desk-22(8)/WL/Research/CR-37(21-22)/1344/21-21, Nagpur, dated 07/09/2021), Telangana (No.26803/2012/WL-2, dated 14/09/2021), Assam (No. WL/FG/31/Research.T.C./28^th^T.C.2021, dated 20/12/2021), Tamil Nadu (No.4822/2021/WL1, dated 28/01/2022), Madhya Pradesh (No./M.H.-II/ Research/2824, Bhopal, dated 13/04/2022), Karnataka (No.PCCF(WL)/E2/CR-37/2021-22, dated 17/05/2022), and Uttar Pradesh (No. 23-2-12(G), Lucknow, dated 20/05/2022).

## Consent for Publication

All authors in our study (Wasimuddin, Gulafsha Khan, Sneha Narayan, Neha Sharma, Basavaraj S. Holeyachi, Md. Abdul Hakeem, Archana Bharadwaj Siva and P. Anuradha Reddy) agree to the publication.

## Data Availability

The data generated and analysed in the current study are available from the corresponding author on reasonable request.

## CRediT Authorship contribution statement

**Wasimuddin** – conceptualization, formal analysis, methodology, software, visualization, writing original draft; **Gulafsha Khan** – sample collection, methodology, investigation, formal analysis; **Sneha Narayan** – sample collection, methodology, data curation, validation; **Neha Sharma** – sample collection, methodology; **Basavaraj S. Holeyachi** – resources, data curation; **Md. Abdul Hakeem** – resources, data curation; **Archana Bharadwaj Siva** – fund acquisition, project administration, resources, supervision; and **P. Anuradha Reddy** – fund acquisition, project administration, supervision, resources, visualization, writing original draft. All authors reviewed and edited the manuscript.

## Acknowledgements

We sincerely thank the Chief Wildlife Wardens of the states of Uttar Pradesh, West Bengal, Assam, Madhya Pradesh, Maharashtra, Telangana, Tamil Nadu and Karnataka for permitting us to collect samples of animals in captivity. We are grateful to the support extended by staff of zoos and rescue centers.

